# Identifying eukaryotes in drinking water metagenomes and factors influencing their biogeography

**DOI:** 10.1101/2022.11.29.518372

**Authors:** Marco Gabrielli, Zihan Dai, Vincent Delafont, Peer Timmers, Paul van der Wielen, Manuela Antonelli, Ameet Pinto

## Abstract

The biogeography of eukaryotes in drinking water systems is poorly understood relative to prokaryotes or viruses. A common challenge with studying complex eukaryotic communities from natural and engineered systems is that the metagenomic analysis workflows are currently not as mature as those that focus on prokaryotes or even viruses. In this study, we benchmarked different strategies to recover eukaryotic sequences and genomes from metagenomic data and applied the best-performing workflow to explore eukaryotic communities present in drinking water distribution systems (DWDSs). We developed an ensemble approach that exploits k-mer and reference-based strategies to improve eukaryotic sequence identification from metagenomes and identified MetaBAT2 as the best performing binning approach for clustering of eukaryotic sequences. Applying this workflow on the DWDSs metagenomes showed that eukaryotic sequences typically constituted a small proportion (i.e., <1%) of the overall metagenomic data. Eukaryotic sequences showed higher relative abundances in surface water-fed and chlorine disinfected systems. Further, the alpha and beta-diversity of eukaryotes were correlated with prokaryotic and viral communities. Finally, a co-occurrence analysis highlighted clusters of eukaryotes whose presence and abundance in DWDSs is affected by disinfection strategies, climate conditions, and source water types.

**Synopsis:** After benchmarking tools and developing a dedicated consensus workflow for eukaryotic sequence detection in metagenomes, the experimental, environmental, and engineering factors affecting their biogeography in drinking water distribution systems were investigated

**Graphical abstract:** 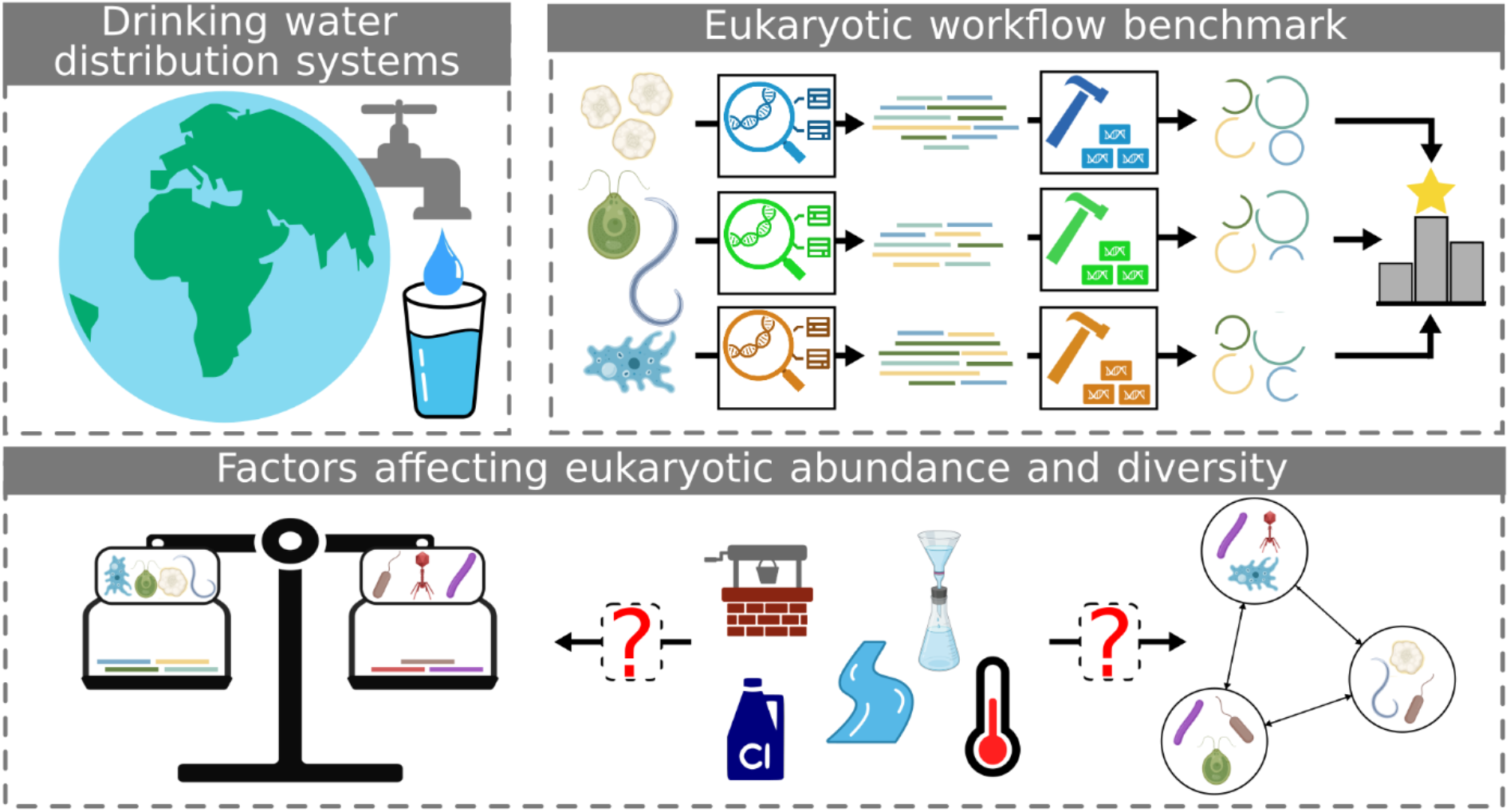

## 1. Introduction

Several drinking water regulations (e.g., ^1–3^) include parasitic eukaryotes such as *Giardia lamblia* and *Cryptosporidium spp*. among the microbial parameters of interest due to their potential negative effects on human health^4,5^. However, compared to prokaryotes and especially bacteria, only few studies have focused on the presence and the ecological role of unicellular and multicellular eukaryotes within drinking water distribution systems (DWDSs). These studies, employing targeted approaches (e.g., internal transcribed spacer – ITS, 18S rRNA) and traditional culture-based methods, showed that variations in eukaryotic community in drinking water systems are associated with water quality characteristics (e.g., organic carbon, nutrients), source water type and disinfection strategies^6,7^. Further, they have shown that several eukaryotes can also shield opportunistic pathogens from disinfection and affect biofilm properties through grazing^8^ and colonization^9^.

Gene targeted approaches (i.e., amplicon sequencing) have been recently used to not only detect a vast diversity of eukaryotes in drinking water systems, but also to probe their activity in different conditions^7^. These techniques have also revealed the widespread presence of eukaryotic pathogens in DWDSs even with the presence of a disinfectant residual^10,11^. However, amplicon sequencing approaches provide limited ecological and physiological insights since they may not permit fine-scale taxonomic resolution and they do not provide any information on their potential functional traits^12^. Furthermore, such insights could also be impacted by several biases, arising from polymerase chain reaction (PCR) amplification, poor comparability of results between different hypervariable regions and variable small subunit (SSU) rRNA gene copy numbers^13,14^.

In contrast, shotgun DNA sequencing (i.e., metagenomics) alleviate these limitations by direct sequencing of extracted genetic material collected in the sample and allowing genome and metabolic reconstructions of detected microorganism^15^; this comes with the drawback of reduced detection limits as compared to amplicon sequencing approaches^16^. Further, metagenomic approaches can prove challenging when dealing with complex genomes such as the eukaryotic ones, especially in case of organisms with low relative abundances^17,18^. In addition, eukaryotic-focused data analysis workflows are relatively less developed in comparison with prokaryotic or viral ones and no extensive comparison among the different options has been conducted.

To this end, this study (i) benchmarked several approaches for eukaryotic sequence detection using synthetic metagenome constructs and then (ii) applied the optimized eukaryotic sequence detection workflow to publicly available DWDS metagenomes to characterize the diversity and biogeography of eukaryotic communities in drinking water. Specifically, we investigate (iii) experimental, environmental, and management factors that may impact eukaryotic detection and (iv) their associations between eukaryotes, prokaryotes, and viruses.

## 2. Materials and methods

### 2.1 Bioinformatics tools benchmarking

#### 2.1.1 Data sources and in silico mock metagenomes construction

The eukaryotic and prokaryotic genomes used to benchmark bioinformatics tools were downloaded from NCBI Genbank^19^, RefSeq^20^, and JGI Genome Portal^21^. A dataset was created including 33 eukaryotic genomes and 216 prokaryotic ones (Table S1). These genomes were selected after determining their absence from training sets of the k-mer based tools tested in this study. To evaluate eukaryotic sequence identification tools, test contigs sets were created by extracting 100 randomly selected sequences of length 1, 3 and 5 kbp from contigs present in the downloaded genomes, similarly to previous studies^22–25^. Benchmark samples and assemblies for eukaryotic sequence binning were obtained using CAMISIM v1.3^26^. Specifically, fifteen mock metagenomes were simulated using the genomes from Table S1, followed by generation of three metagenomic assemblies (five mock metagenomes per assembly). While all parameters were kept as default, the composition of relative genomes abundances in the different samples were drawn from a lognormal distribution (μ = 1, σ = 2) imposing a ratio of total base pairs (bp) equal to 0.05 between prokaryotic and eukaryotic reads in all samples (Table S2).

#### 2.1.2 Benchmarking workflows for the identification of eukaryotic sequences using *in silico* mock metagenomes

EukRep v0.6.6^25^, Tiara v1.0.2^23^, Whokaryote v0.0.1^24^ and DeepMicrobeFinder^22^ were used to identify eukaryotic contigs in the generated test contigs sets using k-mer-based approaches. EukRep and Tiara were implemented using three different thresholds varying from lenient to stricter classifications. Furthermore, majority voting identification (ties excluded) was obtained by combining the results from EukRep, Tiara and Whokaryote, and, alternatively, Tiara, Whokaryote and DeepMicrobeFinder to test the complementarity of the different tools. Sequences were also classified using Kaiju v1.8.2^27^ (nr_euk database version: 2021-02-24) and CAT v5.2.3^28^ (database version: 20210107) which relies on Prodigal^29^ and DIAMOND^30^. Finally, a hybrid strategy combining the results of reference and k-mer-based approaches was tested. This strategy identified eukaryotic contigs using reference-based tools and then used k-mer-based characterization for contigs with no available reference-based annotations. After the classification of each contig, the sequences were randomly subsampled to achieve a eukaryotic to prokaryotic bp ratio equal to 0.05, considered as representative ratios of the two superkingdoms abundances in DWDS metagenomes^31^ to obtain performance estimates representative of real-world situations^22^. To properly account for both false positive and negatives in imbalanced datasets^32^, classification performances were evaluated based on the Matthews Correlation Coefficient (MCC), precision, and recall using yardstick v1.1.0^33^. The subsampling was repeated 100 times, estimating mean and standard deviation of each performance metric. A flow chart of the eukaryotic contigs identification benchmarking is shown in Figure S1.

#### 2.1.3 Benchmarking workflows for binning of eukaryotic sequences using *in silico* mock metagenomes

The gold standard assemblies generated by CAMISIM were classified using a hybrid reference and k-mer-based strategy, imposing a minimum contig length for reference-based and k-mer-based classifications equal to 1 and 3 kbp based on findings from classifier testing (see results in Section 3.1.1), respectively. Contigs were binned with CONCOCT v1.1.0^34^, MetaBAT2 v2.15^35^, SemiBin v0.7.0^36^ and VAMB v3.0.2^37^ according to the following strategies: (i) binning the full assemblies (FULL), (ii) binning only the contigs classified as eukaryotic (EUK-only) or (iii) binning contigs classified as eukaryotic and unclassified contigs (OTHER-rem) (i.e., removing contigs classified as prokaryotes or viruses). To include eukaryotic taxonomic information in SemiBin, contigs taxonomic assignment was performed using CAT. While all the binners were tested using default settings, MetaBAT2 was also run with the parameter minCV equal to 0.1 and 0.33. Each strategy was tested with minimum contig length cutoff of 1, 1.5, and 3 kbp. The binning quality was evaluated focusing on the bins with the majority of the bp derived from eukaryotic genomes. Binning results were evaluated using AMBER v2.0.3^38^ on: (i) the percentage of eukaryotic bp binned, (ii) the trade-off between bins purity and completeness, assessed through the F1 score, and (iii) the similarity between the recovered bins and the original eukaryotic genomes, measured through the Adjusted Rand Index (ARI). A flow chart of the procedure is presented in Figure S2.

### 2.2 Analysis of publicly available DWDS metagenomes

#### 2.2.1 Datasets used

Metagenomes derived from DWDSs were downloaded from NCBI using the SRA Toolkit v2.9.6^39^ or, in case deposited on MG-RAST^40^ retrieved directly from corresponding researchers for a total of 181 distinct samples. Only sequences derived from finished water at drinking water treatment plant (DWTP) and samples from DWDS were considered, excluding the ones either collected in raw water, within drinking water treatment plants and of DWDSs biofilms. Table S3 includes a list of the samples with the details regarding experimental procedures used for each sample^31,41–52^.

#### 2.2.2 Bioinformatic analyses

Raw reads from metagenomes were quality filtered and trimmed using fastp v0.20.1/v0.23.2^53^, followed by vector contamination removal using the UniVec_Core database^19^ and BWA-MEM v0.7.17 or BWA-MEM2 v2.2.1^54^, SAMtools v1.9^55^ and bedtools v2.30.0^56^. Cleaned reads were merged in case obtained in multiple sequencing of the sample and used to estimate the metagenome sequencing coverage using Nonpareil v3.4.1^57^ and screened to identify the number of read pairs properly mapping to the 16S or 18S rRNA genes contained in the SILVA database v138.1^58^ using PhyloFlash v3.4^59^ and SAMtools.

Cleaned reads from samples collected within each distribution system were co-assembled using MetaSpades v3.10.1/v3.15.3^60^ after filtering for contigs greater than 1 kbp using SeqKit v2.1.0^61^. The coverage and depth of retained contigs were determined using BWA-MEM2 and SAMtools. For subsequent analyses, retained contigs were considered present within a sample if at least 25% of the bases in the contigs had at least one read mapping to them. Eukaryotic contigs in metagenome assemblies were identified using a hybrid reference and k-mer-based approach using contigs with minimum contig length of 1 and 3 kbp, respectively. The fractions of eukaryotic, prokaryotic, viral, and unclassified contigs within a metagenome were estimated from the coverage information previously estimated. To capture the diversity within each group, dissimilarity between the contigs present in each sample were estimated using Mash v2.3^62^ using 20,000 randomly sampled contigs within each sample. For each group, MASH was also used to estimate the beta diversity across DWDSs. Before the beta diversity estimation, assemblies with less than 250 contigs were removed, rarefying the remaining assemblies to an equal number of contigs. The existence of significant presence/absence co-occurrence patterns of 18S rRNA genes was evaluated using CoNet v1.1.1^63^ and Cytoscape v3.9.1^64^ using a hypergeometric distribution-based approach and a significance threshold of 0.05. The obtained network was divided in modules maximizing modularity using the Leiden algorithm^65^ implemented in leidenbase v0.1.12^66^. The 18S rRNA gene sequence percentage identity of genes within in each network module was estimated using blastn as carried out by Wu and co-workers^67^.

#### 2.2.3 Statistical analyses

Statistical analyses were carried out in R v4.2.1^68^. Alpha diversity analyses of SSU rRNA genes were performed using breakaway v4.7.6^69^ and DivNet v0.4.0^70^ modelling the effect on samples due to all the categorical factors (i.e., DWDS of origin and abundance cluster membership), while the correlations among eukaryotic and prokaryotic or viral beta diversities were tested using a Mantel test, as implemented in vegan v2.6-2^71^. Linear mixed effects models from the lme4 v1.1-29 package^72^ were used to evaluate the differences in water quality characteristics among different clusters, using a random effect to account for the differences among DWDS. Log transformations were used to correct for residuals’ heteroscedasticity. For each rRNA genes module identified with the network analysis, hurdle negative binomial models in the package countreg v0.2-1^73^ were used to model the number of 18S rRNA genes occurrences of each module in each sample as a function of the disinfection strategy, the source water origin, and the Koppen climate zone^74^.

## 3. Results and discussion

### 3.1 Bioinformatic tools benchmarking

#### 3.1.1 Eukaryotic identification

The majority of the sequenced data in metagenomic assemblies from complex environmental samples are typically contained in short contigs (e.g., < 5 kbp), especially in case of complex communities with low abundance organisms^17,75,76^. However, eukaryotic sequence identification tool benchmarks often focus predominantly on longer contigs^22–25^, potentially leading to an overestimation of the tools’ performances in complex metagenomes. Eukaryotic sequence identification from metagenome assemblies utilized either k-mer signature differences between eukaryotes and prokaryotes, or comparison of unknown sequences with reference databases. As described in previous studies, the performance of k-mer-based strategies improves with increasing contig length (Figure 1a). In our benchmark, EukRep resulted in poorer performances compared to the other tools due to the very liberal eukaryotic classification regardless of the settings used, which is consistent with previous results^22–24^. While this may ensure the recovery of most eukaryotic sequences^25^, this also might result in higher contamination if a thorough contamination removal step is not performed. Instead, in contrast with recent reports^24^, Tiara outperformed Whokaryote. This is due to the fact that the distributions of the gene structure metrics used by Whokaryote depend on contigs length and they are, thus, not generalizable (Figure S3). In fact, while the inclusion of such metrics alongside Tiara’s predictions, as done by Whokaryote, leads to more accurate classifications of long contigs^24^, their inclusion with short contigs is not effective likely due to the presence of incomplete and fragmented genes.

**Figure 1.**
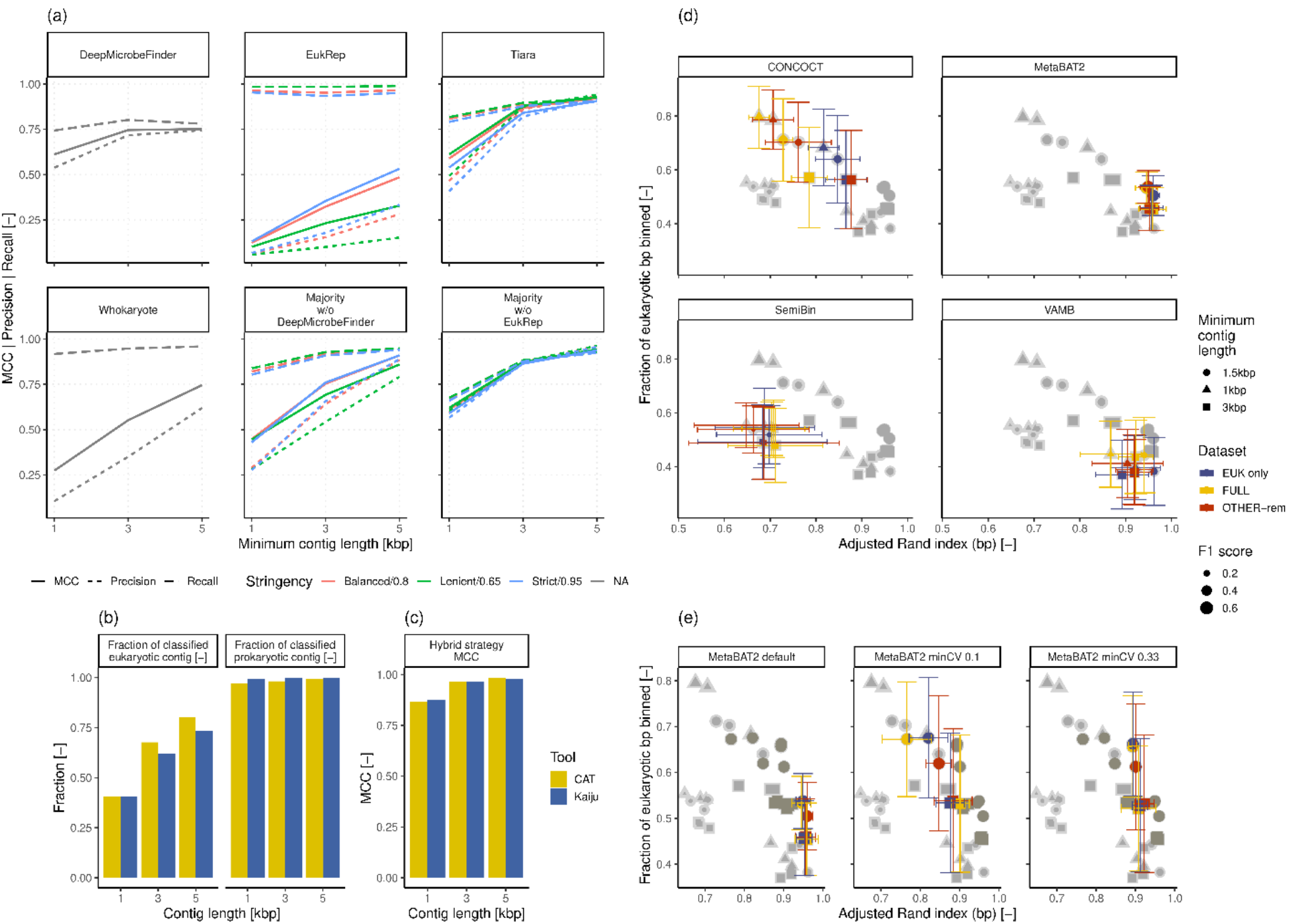
(a) MCC, precision, and recall of tested k-mer-based strategies for eukaryotic identification. The stringency levels tested refer to EukRep’s setting and Tiara’s probability threshold used, while NA indicates tools where stringency was not adjustable. (b) Fraction of eukaryotic and prokaryotic contigs identified by reference-based tools and eukaryotic identification MCC. (c) Eukaryotic identification MCC of hybrid k-mer- and reference-based strategies. (d) Average and standard deviation of the fraction of eukaryotic bp binned and ARI obtained by the tested binners on the simulated metagenomes. (e) Effect of the variation of the MetaBAT2’s parameter minCV on the percentage of eukaryotic bp binned and ARI. In both (d) and (e) the size of the outer marker depicted in light grey represents the F1 score estimated using only the most complete bin per each eukaryotic genome recovered, while the size of inner marker represents the value including all generated bins. For both subplots the colored symbols highlight the markers related to binner or setting stated in the facet title, while greyed-out markers relate to the other binners/settings to aid comparison between the results.

Compared to the other tools, DeepMicrobeFinder was trained on short contigs (i.e., 0.5 – 5 kbp)^22^. This resulted in relatively high MCC values at 1kbp but led to only a limited improvement with longer contigs, reaching a plateau of its MCC value at 3 kbp (Figure 1a). As the various tools present different strengths and weaknesses^24^, we tested the performance of majority voting strategies (ties excluded) using either the combination of EukRep, Whokaryote and Tiara (Majority w/o DeepMicrobeFinder) or DeepMicrobeFinder, Whokaryote and Tiara (Majority w/o EukRep) against Tiara, the best-performing single tool. The inclusion of EukRep alongside Tiara and Whokaryote (i.e., Majority w/o DeepMicrobeFinder) resulted in lower performances compared to Tiara, due to the low precision of both EukRep and Whokaryote. Instead, the use of DeepMicrobeFinder, Tiara and Whokaryote (i.e., Majority w/o EukRep), resulted in MCC values approximately 3% greater than Tiara, due to an increase in precision obtained at the cost of a drop in recall. Noticeably, the larger MCC improvement between 3 and 5 kbp obtained by Majority w/o EukRep (8%) compared to Tiara (6%) suggests further improvements over Tiara with longer contigs.

In addition to k-mer-based tools, two reference-based tools were tested (Figure 1b). In contrast to k-mer based approaches, both Kaiju and CAT presented MCC values above 0.99 for 1-kbp-long contigs, in concordance with previous benchmarks^28^. However, this high MCC was associated with the loss of large fractions of eukaryotic contigs which were not classified, especially for shorter contigs; this is likely because of the presence of incomplete fragmented genes^77^. We further tested the integration of the two approaches to combine the ability to classify all sequences of k-mer-based tools with the high-accuracy of reference-based approaches. Such hybrid strategy classifies contigs primarily using reference-based predictions, resorting to k-mer-based results in case no reference-based annotation is available. The integration of reference-based strategies with Majority w/o EukRep improved overall performance compared to the use of exclusively reference-or k-mer-based strategies, regardless of the reference-based tool used (Figure 1c). In fact, CAT provided a 0.5% higher MCC value than Kaiju with 5 kbp-long contigs, while Kaiju resulted in a 1.3% higher MCC value with 1 kbp contigs. This ensemble approach for identification of eukaryotic sequences from metagenomic data is documented as a workflow, EUKsemble (https://github.com/mgabriell1/EUKsemble). This workflow combines the results of Majority w/o EukRep with Kaiju’s or CAT’s prediction to improve eukaryotic sequence retrieval from metagenomic assemblies. As the combination between k-mer-based strategies and CAT or Kaiju leads to similar results, the choice between the two is left to the user, allowing the use of Kaiju in case available computing resources are limited^28^. Noticeably, to maximize eukaryotic retrieval while minimizing false positives, different minimum contig lengths for the two classification strategies can be exploited. For instance, the chosen reference-based tool could be applied to very short contig lengths (e.g., 1 kbp) in order to retrieve with high confidence as many eukaryotic contigs as possible, while Majority w/o EukRep could be used with contigs longer than 3 kbp, length at which such strategy provides sufficient performances.

#### 3.1.2 Recovering eukaryotic metagenome assembled genomes from metagenomic assemblies

Previously published eukaryotic-targeted metagenome pipelines rely prevalently on MetaBAT2 or CONCOCT binning either directly on the whole metagenome or eukaryotic-screened contigs^25,78,79^. However, currently no direct comparison between the various alternatives is present. Hence, we tested the performance of several state-of-the-art binners either after the selection of eukaryotic contigs (EUK-only) or directly on the whole metagenome (FULL) using a range of minimum contig lengths. Furthermore, as eukaryotic identification was performed using EUKsemble using different minimum contig length for k-mer- and Kaiju-based identification (i.e., 3 and 1 kbp), we also tested the possibility of performing binning with contigs identified as Eukaryotes or without any assigned superkingdom (OTHER-rem) after the removal of only the contigs classified as non-eukaryotic (i.e., prokaryotic, viral).

Irrespective of the tool used, binning suffers inherently from a tradeoff between the amount of assembled bp included in the recovered bins and the binning quality (Figure 1d), as reported by similar benchmarks^80^. Our results indicate that this is observable both across different tools, minimum contig lengths, and binning strategies (Figure 1d). CONCOCT generally recruits the most eukaryotic bp into bins, while conversely MetaBAT2 and VAMB maximize the ARI, indicating higher quality of the reconstructed eukaryotic bins. SemiBin performs poorly on both metrics. Increasing the minimum contig length thresholds, on one hand, improves the bin quality for all binning tools (i.e., higher ARI values), while coincidentally results in lower fractions of eukaryotic bp within bins. Direct binning of the entire metagenome allows recovery of a higher fraction of eukaryotic bp present in the metagenome, but at the cost of a lower ARI as compared to binning exclusively on contigs annotated as eukaryotic. The limited eukaryotic recovery arises due to the limits of reference-based eukaryotic identification which do not classify a large fraction of the eukaryotic contigs between 1 and 3 kbp (Figure 1b). In contrast, excluding only the contigs classified as non-eukaryotic provided an increase in the ARI value for all binners except for SemiBin and especially using CONCOCT (average percentage increase: CONCOCT: 7%, MetaBAT2: 1%, VAMB: 1%) compared to binning the entire metagenome, while recovering up to 15% more bp than binning exclusively contigs annotated as eukaryotic. While the selection between binning only the contigs identified as eukaryotic or including also the non-classified ones can depend on the desired levels of purity of the recovered bins and may require extensive curation (e.g., Delmont and colleagues^81^), this practice can result in highly chimeric bins including both eukaryotic and prokaryotic contigs (Figure S4) which are likely to affect downstream results.

As MetaBAT2 (with default settings) resulted in the highest ARI, we tested whether it was possible to increase the recovery of eukaryotic data by including contigs with low coverage depth. Indeed, as shown in Figure 1e, reducing the value of minCV, the parameter controlling the minimum coverage depth admissible, increased the recovery, reaching values comparable to CONCOCT. However, excessively low minCV values (i.e., 0.1) affected ARI values negatively, indicating the value of 0.33 as a suitable lower bound. The need to adjust this parameter does not require prior knowledge on the microbial community analyzed, but only a previous identification of eukaryotic contigs and coverage depth estimation. The highest average F1 scores of the most complete bin per recovered genome were provided by MetaBAT2 with reduced minCV parameter values and CONCOCT, followed by default MetaBAT2, SemiBin and VAMB, mostly due to variations in completeness, as purity showed high average values (i.e., > 0.9) (Table S4). However, when considering all the recovered bins, MetaBAT2 led to the highest scores since the other binners show high fragmentation of the initial genomes. This fragmentation was not associated with chromosomal organization of eukaryotic genomes, but rather due to the combination of high k-mer diversity and low coverage depth (Figure S5, Meyer and colleagues^17^). Both the eukaryotic identification and binning benchmark results highlight how contigs length plays a significant role in eukaryotic recovery from metagenomes. Considering these results, metaSPAdes co-assembly appears to be the most appropriate assembler for eukaryotes as known to produce longer contigs than other assemblers and single-sample assembly strategies^17,76,82^.

### 3.2 Factors affecting eukaryotic abundance in DWDS metagenomes

The eukaryotic fraction of DWDS metagenomes was assessed on a total of 181 samples collected from 81 DWDSs across the globe (Figure 2a) using EUKsemble relying on Kaiju’s reference-based approach. Despite the intensive eukaryotic identification procedure, even though few studies showed particularly high fractions of bp mapping to eukaryotic contigs, most DWDS showed fractions of eukaryotic bp below 1% and in most cases lower than the fraction mapping to viral contigs, identified based on Kaiju’s classification. Such value is similar to previous results^31,51,83^ and is correlated with the relative abundance of eukaryotic 18S rRNA genes detected from the metagenome data (Figure S6). Still, the retrieved eukaryotic percentages are likely underestimated due to the exclusion of very short contigs (< 1 kbp) and the limits of eukaryotic identification (see Figure 1b). Indeed, despite the possible confounding effect caused by the heterogeneity within the data, higher fractions of assembled and identified eukaryotic bp are associated with higher eukaryotic diversity within samples, especially in disinfected system where most data is available (Figure 2b; Spearman correlation disinfected systems: 0.48, p-value: < 0.001; non-disinfected systems: 0.35, p-value: 0.081), further suggesting that eukaryotic populations are systematically under-sampled using current metagenomic approaches; this is in line with the trends for viruses recovered in non-disinfected systems (Figure S7) and in contrast with prokaryotes where the data available showed no significant relationship between their bp fraction and their diversity (Figure S8).

**Figure 2.**
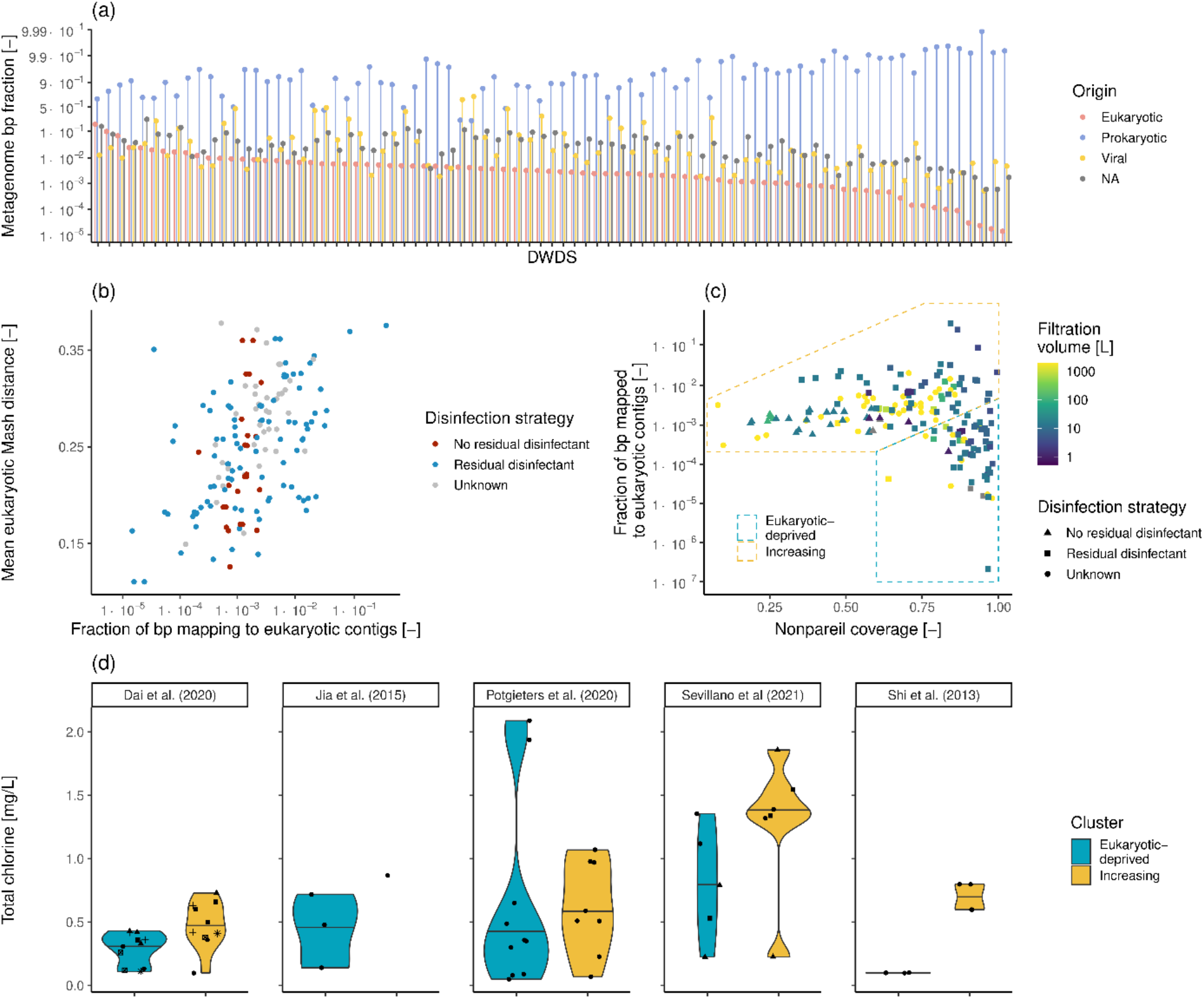
(a) Fractions of eukaryotic, non-eukaryotic and not classified (NA) bp in each of the analyzed metagenomes. EUKsemble results refinement is based on the classification provided by Kaiju. A logit scale was applied to improve clarity of presentation. (b) Association between the fraction of bp mapping to eukaryotic contigs and the mean Mash distance between eukaryotic contigs in each sample as a function of the disinfection strategy employed. (c) Relationship between the Nonpareil coverage and the fraction of bp mapping to eukaryotic contigs in each sample. (d) Total chlorine concentrations from DWDSs whose samples belong to both the eukaryotic-deprived and increasing clusters. The different shapes in each facet in panel (d) indicate different DWDSs.

Differences in the recovery of eukaryotic DNA in metagenomes could be due to different experimental protocols used in the various studies (Table S3), ranging from sample collection to sequencing strategies used. Common filter size ranges for microbial concentration from 0.2 to 0.45 μm and should not affect eukaryotic recovery^84^. The eukaryotic metagenome fractions do not correlate with filtered water volume (p-value = 0.69) reported in corresponding studies (Figure 2c), indicating that filtering a larger volume of water does not improve eukaryotic recovery in metagenomes. Specifically, while filtering larger volumes may increase the number of eukaryotic cells captured, their ratio relative to prokaryotes and viruses will not change and thus may not result in greater recovery of eukaryotic sequences in metagenomes. DNA extraction prior to metagenomic analysis can also affect microbial community recovered using metagenomic sequencing strategies^85,86^. However, as samples taken from the same DWDS were extracted using the same extraction method per DWDS it was not possible to assess the effect of this factor. In any case, while commercial extraction methods were shown to be able to successfully extract eukaryotic DNA^87,88^, specific processing techniques^89^ or the use of dedicated enzymes^90^ might further increase yields and/or quality. Noteworthy, the variety of eukaryotic phenotypes (e.g., soft-shelled, hard-shelled) complicates the ideation of a single optimal extraction method and possibly exacerbates the extraction bias^91^. Lastly, sequencing depth affects the ability to recover rarer taxa^92^. However, increased sequencing efforts did not lead to a significant increase in eukaryotic fractions (p-value = 0.38) due to the confounding effect caused by the different complexity of the microbial community in each sample^93^. Indeed, irrespective of the actual sequencing depth, a better characterization of the microbial community, as indicated by higher Nonpareil coverage^57^, allows higher eukaryotic fractions in metagenomes (Figure 2c), providing evidence of the underestimation of the real presence of eukaryotes in DWDSs.

Despite exhibiting high Nonpareil coverages, some samples show extremely low eukaryotic fractions, separating an “eukaryotic-deprived” cluster from the “increasing” one; this included samples originating from the same DWDS splitting into these two clusters. The two clusters show a significantly different composition with respect to source water type (Chi-Square test, p-value < 0.001), with the eukaryotic-deprived cluster enriched in samples derived from groundwater-fed systems (23%) compared to the increasing cluster (5%), suggesting higher eukaryotic abundances in surface water-fed systems in concordance with what observed in raw waters^94^. However, samples from surface water-fed DWDSs were abundant in both clusters (eukaryotic-deprived cluster = 42.6%, increasing cluster = 29.2%). Water disinfection is an important factor affecting the drinking water microbiome^31,95^. In fact, when comparing the total chlorine concentrations in samples obtained from the same DWDS, but belonging to the different clusters, eukaryotic-deprived samples presented lower chlorine concentrations (95% confidence interval: -195 – -29%, Figure 2d). Eukaryotes are typically have higher resistance to disinfectants compared to bacteria^6,96,97^ and thus, the higher chlorine concentrations present in samples belonging to the increasing cluster could have altered the relative abundances leading to a higher eukaryotic DNA recovery in the metagenomes similar to what observed by Dai and colleagues^31^. In fact, higher chlorine concentrations might limit prokaryotic growth within DWDSs despite the presence of available nutrients^98^ and maintain abundances similar to the ones at the water treatment outlet. On the other hand, several countries limit prokaryotic growth by reducing the nutrients available (i.e., carbon, nitrogen, etc.) in finished drinking water^98^. Indeed, several samples with low Nonpareil coverages from non-disinfected systems are included in the increasing cluster, suggesting limiting microbial growth may be associated with enhance eukaryotic detection in metagenomes. This consideration, coupled with the results presented in Figure 2d, highlights the importance and the interaction of the multiple stresses (i.e., disinfection, nutrient limitation) in shaping drinking water microbiology. Finally, samples belonging to the eukaryotic-deprived cluster present lower average estimated eukaryotic richness (p-value = 0.002) and non-significant changes in Simpson and Shannon diversities (p-values > 0.28) than samples belonging to the increased cluster at similar Nonpareil coverages (i.e., > 0.75), indicating the presence of less diverse and more even communities. Besides water sources and DWDS management strategies, these observations could be associated with other several factors which could not be included in this study due to the missing information. For example, both water treatments, water physico-chemical quality and location within DWDSs have been shown not just to affect prokaryotic, but also eukaryotic abundances^6,99,100^ and should be the focus of targeted studies.

### 3.3 Factor affecting eukaryotic diversity in DWDS metagenomes

As environmental factors and DWDS management strategies affect the proportion of eukaryotes, prokaryotes, and viruses in DWDSs metagenomes, these factors could also affect the taxa present and the diversity across DWDSs. In fact, eukaryotic beta diversity correlates positively with both the prokaryotic and viral ones (Figure 3a, eukaryotic - prokaryotic Mantel statistic r = 0.25, p-value = 0.002; eukaryotic - viral Mantel statistic r = 0.21, p-value = 0.014). While such low values are likely caused by the heterogeneity of upstream treatments and water conditions in the various studies, the presence of correlations among beta diversities suggests that spatiotemporal dynamics and factors which were found to influence prokaryotes and viruses (e.g., disinfection strategies, seasonality, water age)^98^ are likely to be relevant also for eukaryotes. Such concordance is likely the result of both direct causes affecting both eukaryotes and other taxonomic groups (e.g., upstream water treatment, nutrient availability, disinfection stress)^7,99^ or could arise indirectly as a result of their interactions. In fact, depending on environmental stresses (i.e., nutrient availability), fungi have been shown to modulate bacterial growth levels^101^, while protists can both serve as hosts and/or select of specific prokaryotic symbionts and viruses, favoring their multiplication^102–104^, or selectively predate on them^105,106^, highlighting the role of eukaryotes in shaping microbiomes.

**Figure 3.**
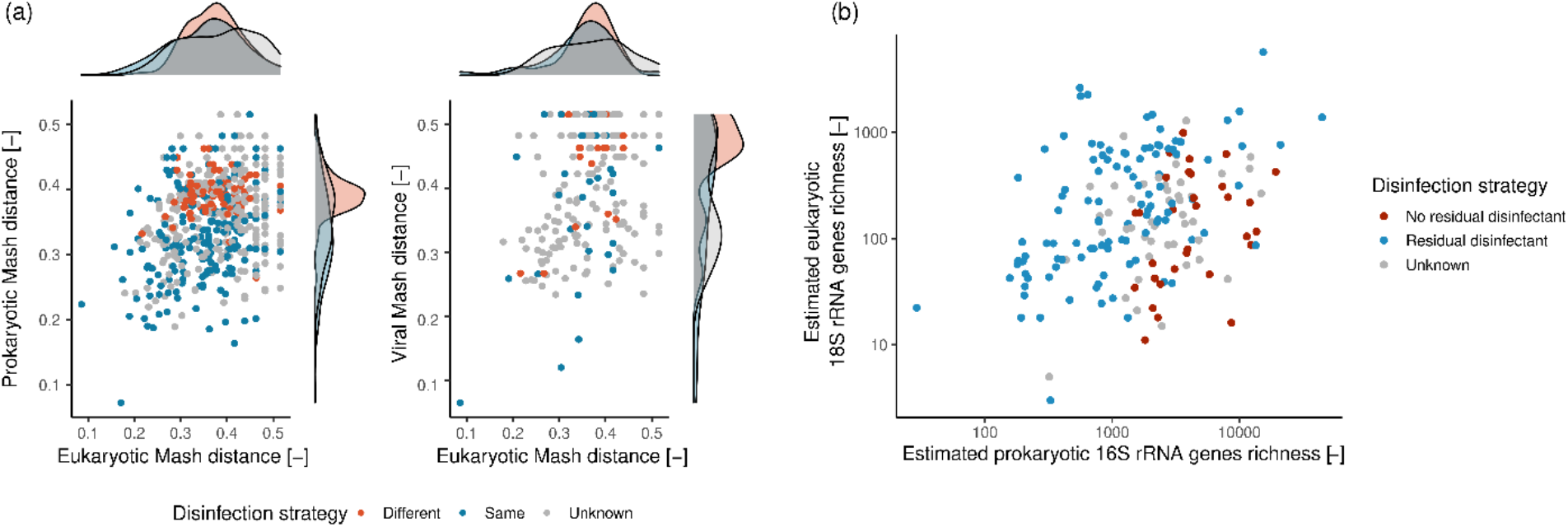
(a) Eukaryotic, prokaryotic and viral beta diversity correlations and marginal distributions as a function of the samples disinfection strategy. (b) Estimated richness of eukaryotic and prokaryotic rRNA genes with respect to disinfection strategy.

Furthermore, through the analysis of the 18S and 16S rRNA genes, it was possible to show a positive correlation between the estimated eukaryotic and prokaryotic richness (disinfected systems: = 0.5, p-vale < 0.001; non-disinfected systems = 0.36, p-value = 0.046). The presence of such correlation, observed also by Yeh and Fuhrman^107^, is concordant with the “diversity begets diversity” hypothesis^108^, likely arising due to interactions between populations across superkingdoms^6^ which expand the availability of ecological niches and thus enhances diversity. Both Figures 3a and 3b, further underline the effect of disinfection of the DWDS microbiome, highlighting, in accordance with to Dai^31^ and Hegarty^109^ and colleagues, the effect of disinfection strategies on the beta diversity of prokaryotic and viral communities in DWDSs and suggesting a lower effect for eukaryotes (median Mash differences: eukaryotes = 0.021; prokaryotes = 0.068; viruses = 0.043). While this result is concordant with the higher chlorine resistance of eukaryotes^6,96,97^, it should be noted that given the likely undersampling of eukaryotic communities in DWDSs, such result might be biased towards the most abundant eukaryotes and that further dedicated studies would be needed to confirm it.

A presence/absence co-occurrence of the 18S rRNA genes present in the samples was carried out to obtain more insights in the eukaryotic communities in the DWDS metagenomes. This approach was favored compared to a relative abundance-based co-occurrence analyses to limit the confounding effects caused by the different experimental protocols employed in the different studies. This network analysis indicated the presence of 11 18S rRNA gene modules, each composed of more than 10 eukaryotic taxa (Figure 4a, Table S5). A 18S rRNA gene sequence similarity analyses within each module indicated that between 13% and 86% of the genes in each module belong to taxa within the same order^67^, with several members belonging to the same family or genus (Figure 4b). The variation in the ranges of percentage identities distributions and the shape of the density distributions suggest the presence in each module of different groups of phylogenetically similar taxa with different degrees of phylogenetic relatedness, possibly arising from several evolutionary and ecological factors^110^. It is important to note that the co-occurrence patterns retrieved here do not necessarily confirm ecological interactions^111^ and should be confirmed by further hypothesis-driven studies. This is especially valid for phagotrophic organisms, while less so for eukaryotes which can feed on other eukaryotes such as nematodes^112^. Noteworthy, nematodes make up most of the nodes in modules 4, 7 and 9. While little information on their diet in drinking water systems is available, some studies in other environmental matrices report that certain species are known to prey on the same, possibly eukaryotic, microorganisms (i.e., fungal-feeder *Aphelenchoides spp*., present in module 4) or even other nematodes (i.e., genus *Mesodorylaimus*, present in module 4), possibly explaining in part of the associations found^112,113^. In addition, some of the co-occurrences retrieved which involve parasitic nematodes are possibly due to the infection of similar hosts, as the plant parasites *Longidorus spp*. and *Xiphinema spp*.^114^ (present both in module 4).

**Figure 4.**
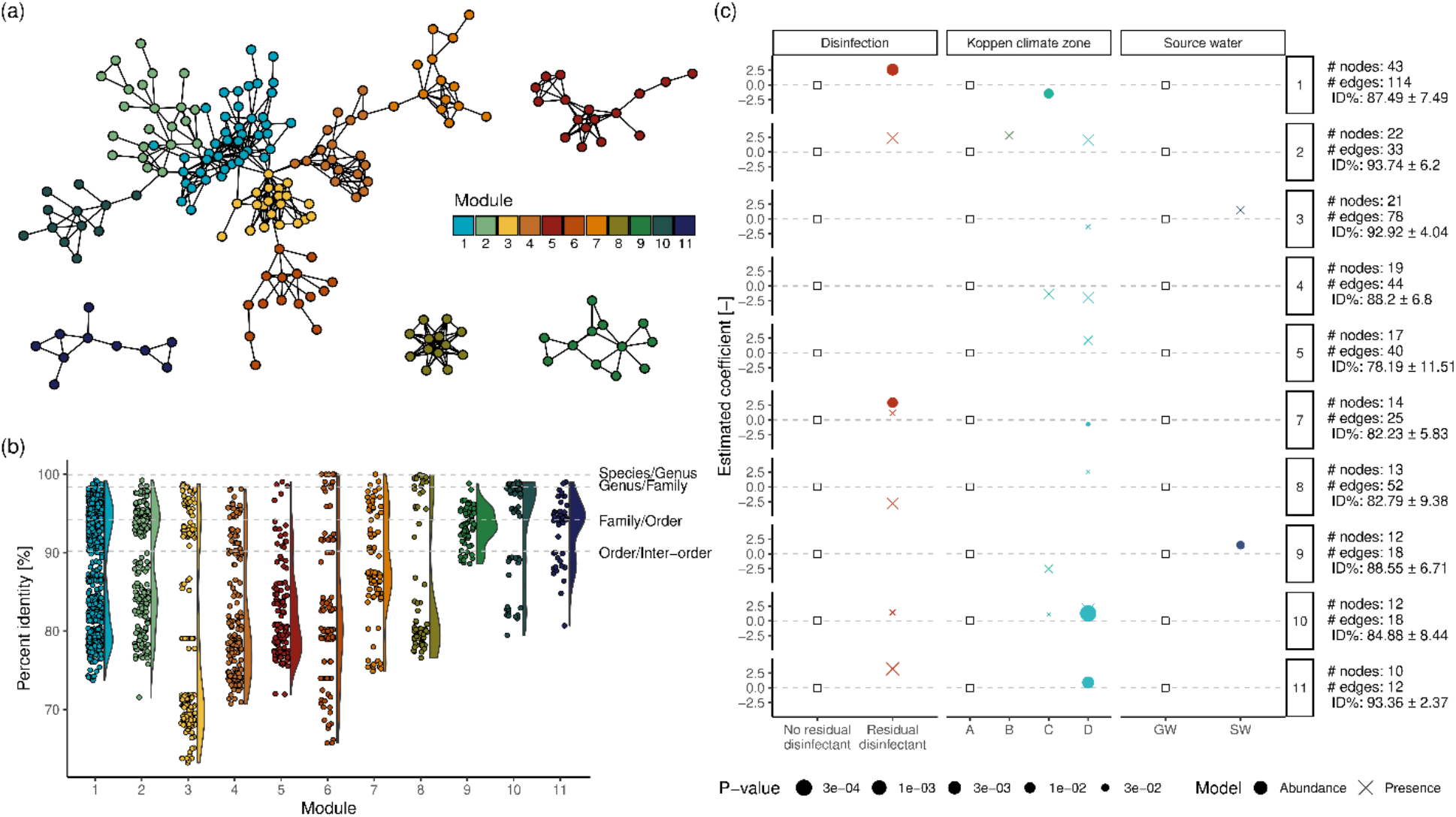
(a) Network of co-occurring eukaryotic 18s rRNA genes colored by module. (b) Percentage identity between 18S rRNA genes belonging to the same module. (c) Estimated coefficients regarding module presence and abundance as a function of disinfection strategy, climate zone and source water type. The coefficients in panel (c) are to be interpreted as relative effects to the factors marked with a white square. To improve clarity the factor “Unknown” was not shown, together with the modules with no significant predictors. Information of the right side of panel (c) reports the number of nodes and edges within each module, and the average and standard deviation of the 18S rRNA genes sequence identities belonging to the module.

The detected association can be considered as groups of eukaryotes which present similar responses to environmental and DWDS management factors or even other (micro)biota. Such interpretation is supported by the analysis, within each module, of the number of occurrences as a function of DWDS disinfection strategy, climate zone, and source water origin. Except for module 6, which did not show any significant predictors (i.e., p-value < 0.05) of its presence or abundance, all the modules showed variations due to the tested factors (Figure 4c). While the higher presence/abundance detected for some modules in disinfected systems might be due to the generically higher Nonpareil coverage of samples derived from such systems, noticeably module 8 shows lower presences in disinfected systems indicating potentially the higher sensitivity of its members to disinfection or the adaptation to low nutrient conditions of non-disinfected DWDSs. In fact, some nodes included in module 8 represent fungi for which some species are known to proliferate under oligotrophic conditions^115^ and demonstrate higher chlorine resistance than viruses and prokaryotes, but lower than protists cyst and oocysts^96,116^. Furthermore, climate zone D (i.e., continental) showed in most cases differences in the presence/abundance of the modules’ members compared to zone A (i.e., tropical), in accordance with the expected climatic differences and geographical distances among the two zones^74^. Finally, in accordance with the results of Section 3.2, higher presence/abundances were observed in drinking water produced from surface water for selected modules. For example, module 3 is composed mostly of *Eustigmatophyceae*, a lineage of photosynthetic algae present in freshwater^117^, indicating the possible role of the eukaryotes (and/or their genetic material) in source waters in seeding downstream DWDSs. Besides the factors taken into the accont in this analysis, it is important to note that several other factors might have affected modules presence/abundance (e.g., upstream treatment, physico-chemical water quality, degree of eukaryotic community characterization).

## 4. Implications for future research and drinking water systems

The results of this study highlight the under-representation of eukaryotes within current DWDS metagenomes. To overcome this issue sampling protocols and extraction methods should be adapted to enrich for eukaryotic microorganisms as already done in different fields surveys, where sampling and laboratory techniques are tailored depending on the microorganisms of interest. For example, laboratory protocols used in the Tara Oceans Expedition were either carefully selected among existing ones or specifically developed in order to limit potential biases and ensure the quality and comparability of the results^89^. Furthermore, this expedition applied a comprehensive sampling strategy which selected for different microorganisms using a size-fractionation approach, based on previously available data^118^. Finally, a wide set of environmental conditions were also collected to aid the data interpretation^119^. In the drinking water field, a similar standardized initiative was carried out by The Netherlands to monitor macroscopic invertebrates using optimized sampling techniques and microscopic techniques^120^, but is not widely adapted.

Despite the recent growing attention eukaryotic-focused metagenomics is not as well established compared to the prokaryotic or viral counterparts, with studies reporting its complexity with current approaches^18^. Compared to prokaryotes and viruses, both the reference data and the option of tools available for eukaryotes is limited, further exacerbating the complexity of the reconstruction of their genomes. In addition, due to the wealth of data provided by marine expeditions, reference databases included in several tools are highly skewed towards marine taxa (e.g., Levy Karin and collaborators^77^, Vaulot and collaborators^121^), potentially limiting and biasing the analyses performed on other environments. While new tools will improve the analysis of eukaryotes from mixed metagenomes, only focused sampling efforts will be able to overcome the compositional bias of current references and enable a clearer view of eukaryotes in DWDSs. In fact, a better understanding of the effects influencing eukaryotic communities, their associations with other microorganisms and viruses would allow to better understand the effect of current microbiological management strategies in DWDSs (e.g., disinfection and nutrient starvation) and devise ecologically-informed microbiological management plans, improving both water treatment and distribution (e.g., Cavallaro and co-workers^122^, Derlon and co-workers^123^).

## Supporting information

Supplementary Informations

## Supporting information

- Supplementary Tables (.xlsx) file provides detailed data regarding the bioinformatic tools benchmarking, the samples used in the analyses and the nodes in the co-occurrence network.
- Supplementary Figures (.docx) file provides additional figures discussed in the main manuscript.

## Acknowledgement

This research was supported by NSF CBET 2220792 and by CAP Holding S.p.A. which funded the PhD grant of Marco Gabrielli.

