## Supplementary Informations for "Identifying eukaryotes in drinking water metagenomes and factors influencing their biogeography"

**Eukaryotic identification in drinking water metagenomes and factors influencing their presence and diversity**

Number of pages: 7; number of tables: 5; number of figures: 8

Table S1. Details of the genomes used for benchmarking eukaryotic identification and binning

Table S2. Distribution of reads in all the simulated samples using CAMISIM

Table S3. Details regarding metagenomic samples used in the analysis

Table S4. Quality metrics of eukaryotic binning on the simulated mock assemblies

Table S5. Eukaryotic 18S rRNA genes belonging to the modules analyzed

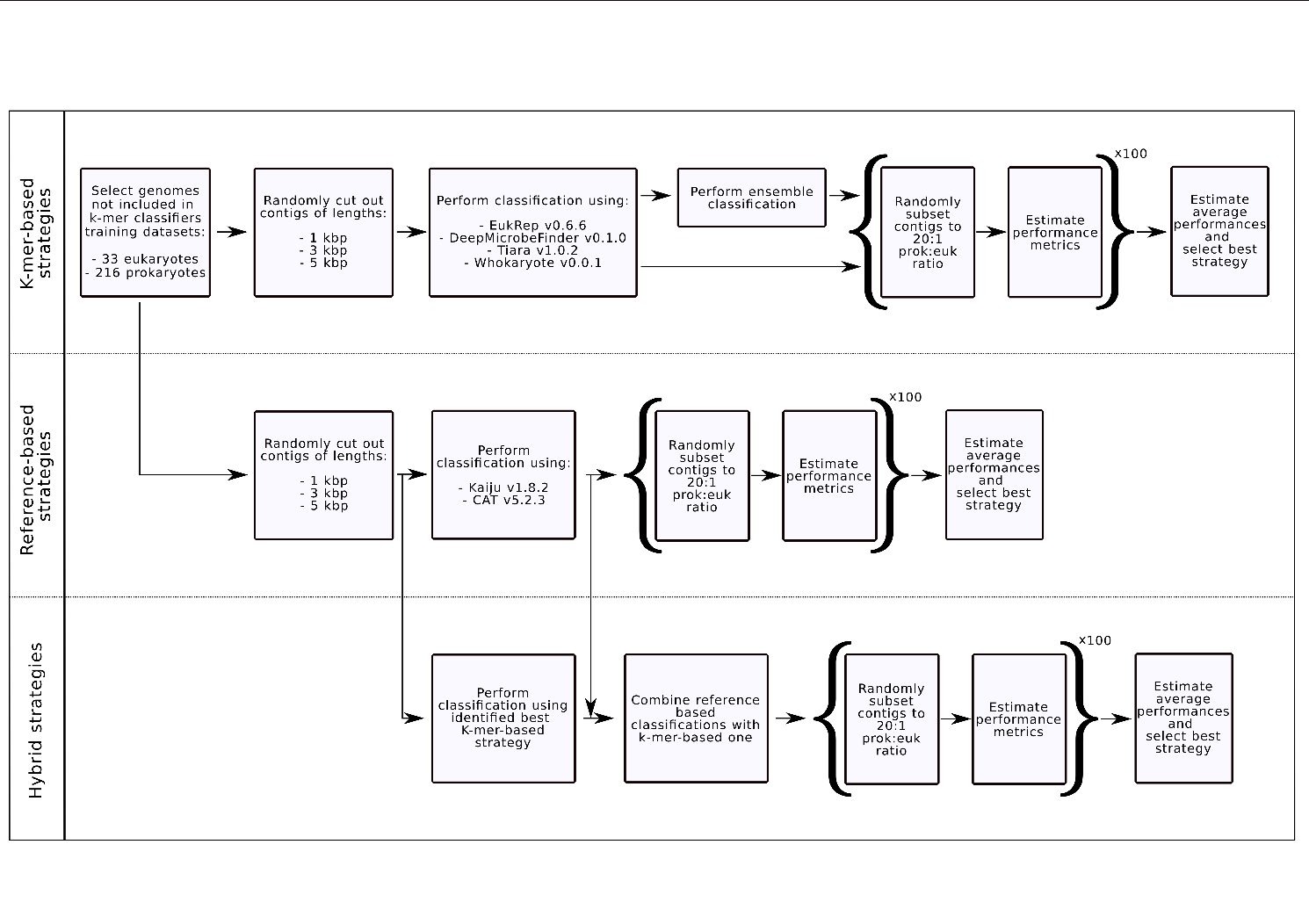
Figure S1. Eukaryotic identification benchmarking workflow for k-mer-based, reference-based and the hybrid strategy. Brackets indicate steps repeated for the number of times indicated (i.e., x100).

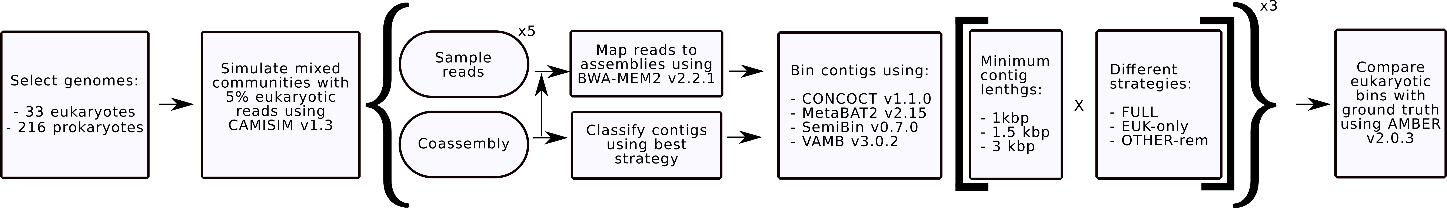
Figure S2. Eukaryotic binning benchmarking workflow. Rounded shapes indicate the files provided by CAMISIM. The square brackets include the combinations of binning strategies tested. Curly brackets indicate steps repeated for the number of times indicated (i.e., x3).

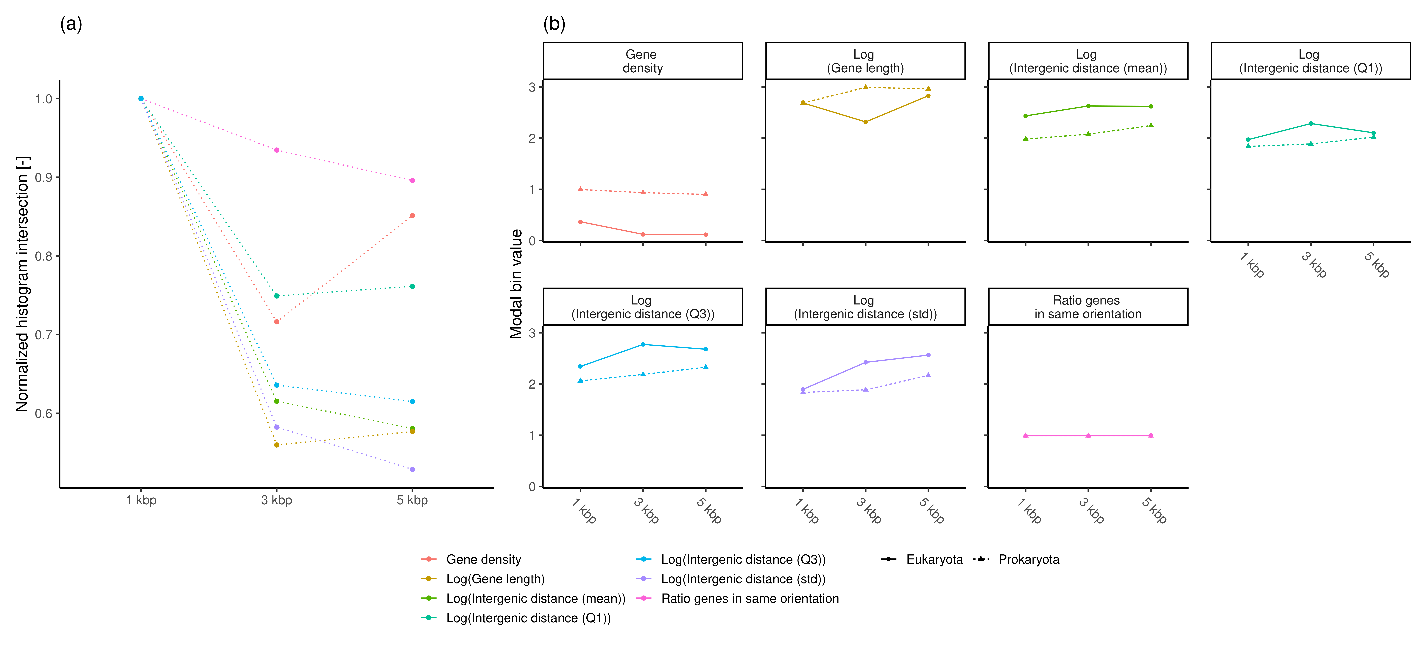
Figure S3. (a) Normalized intersection of the histograms of Whokaryote predictors calculated on eukaryotic and prokaryotic contigs and (b) modal bin values of the histograms of the predictors for eukaryotic and prokaryotic contigs as a function of contigs length.

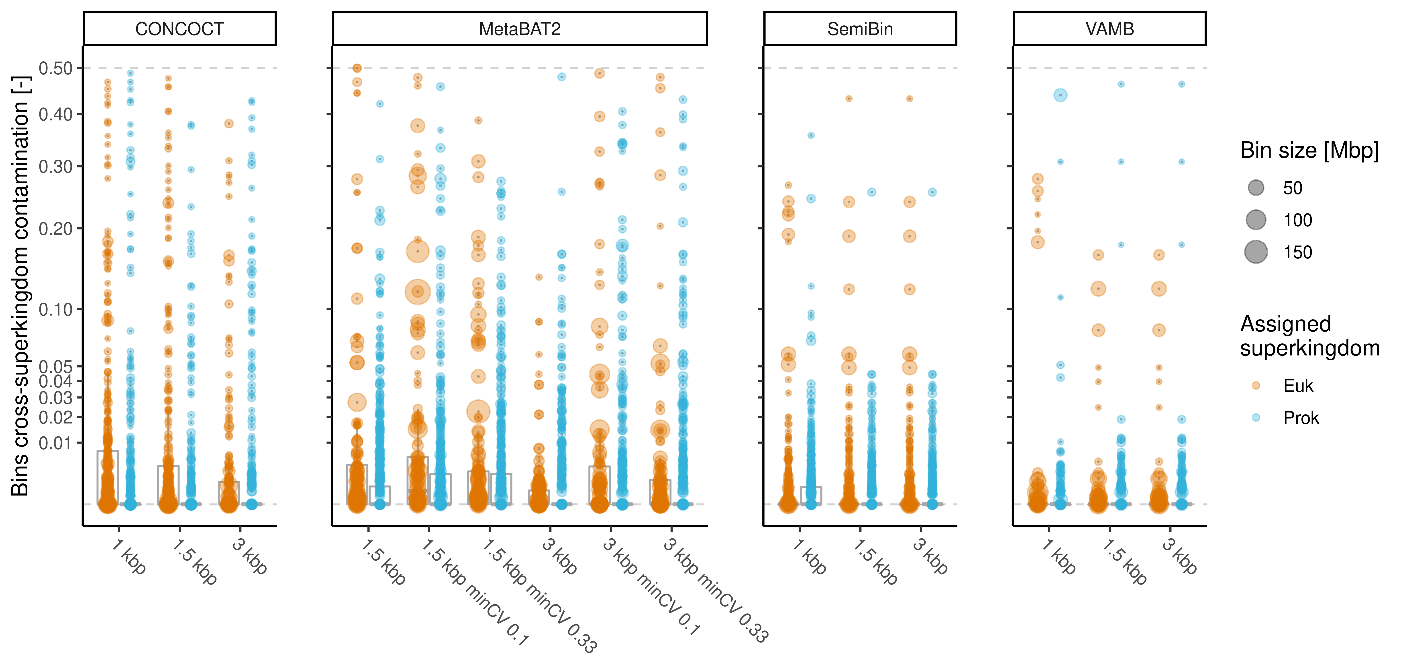

Figure S4. Cross-superkingdom contamination of the bins recovered by the tested binning algorithms. Bins were assigned a superkingdom based on the origin of the majority of the contigs within them. Cross-superkingdom contamination was estimated based on the fraction of contigs length not belonging to the assigned superkingdom with respect to the total length.

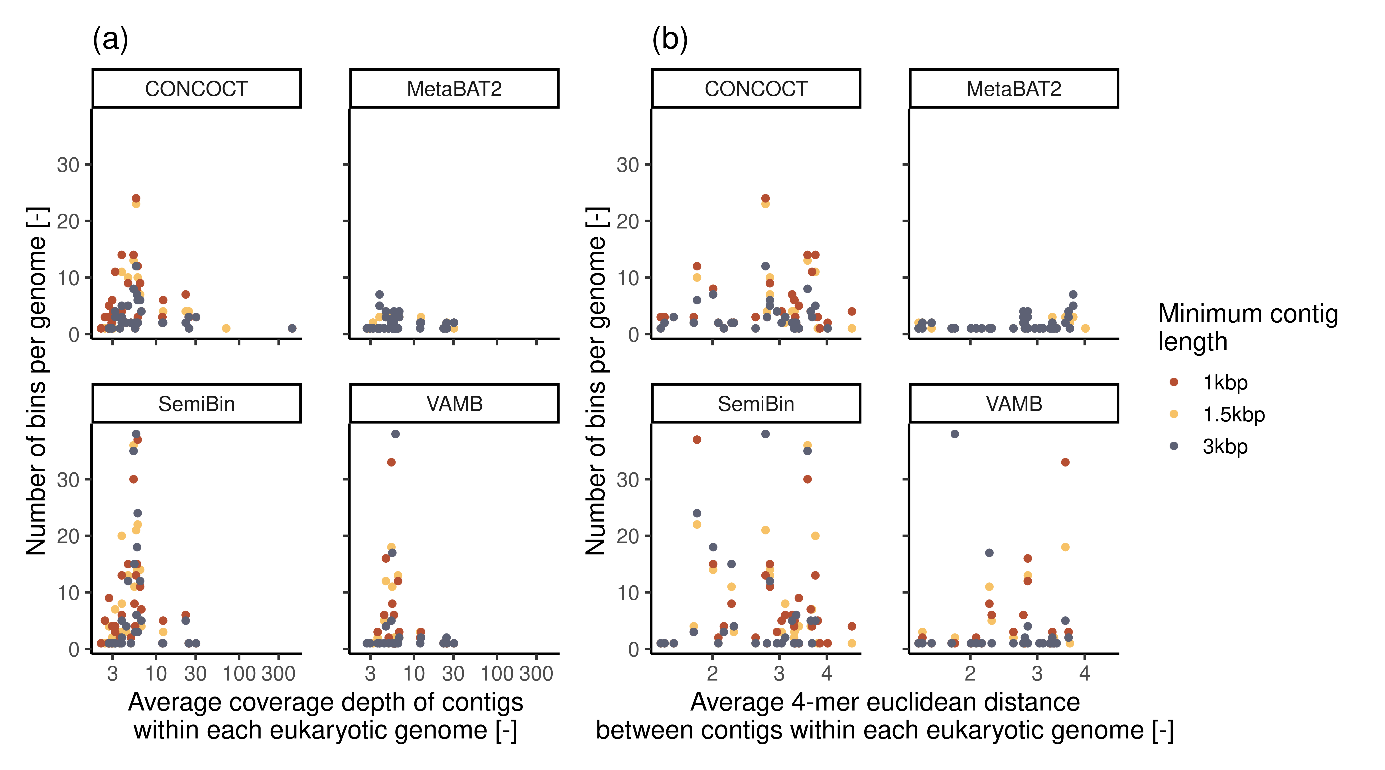

Figure S5. Fragmentation of eukaryotic genomes, shown as the number of bins containing the majority of bp from the same genome, (a) as a function of average contig depth in each genome and (b) the average 4-mer Euclidean distance within the contigs derived from each genome.

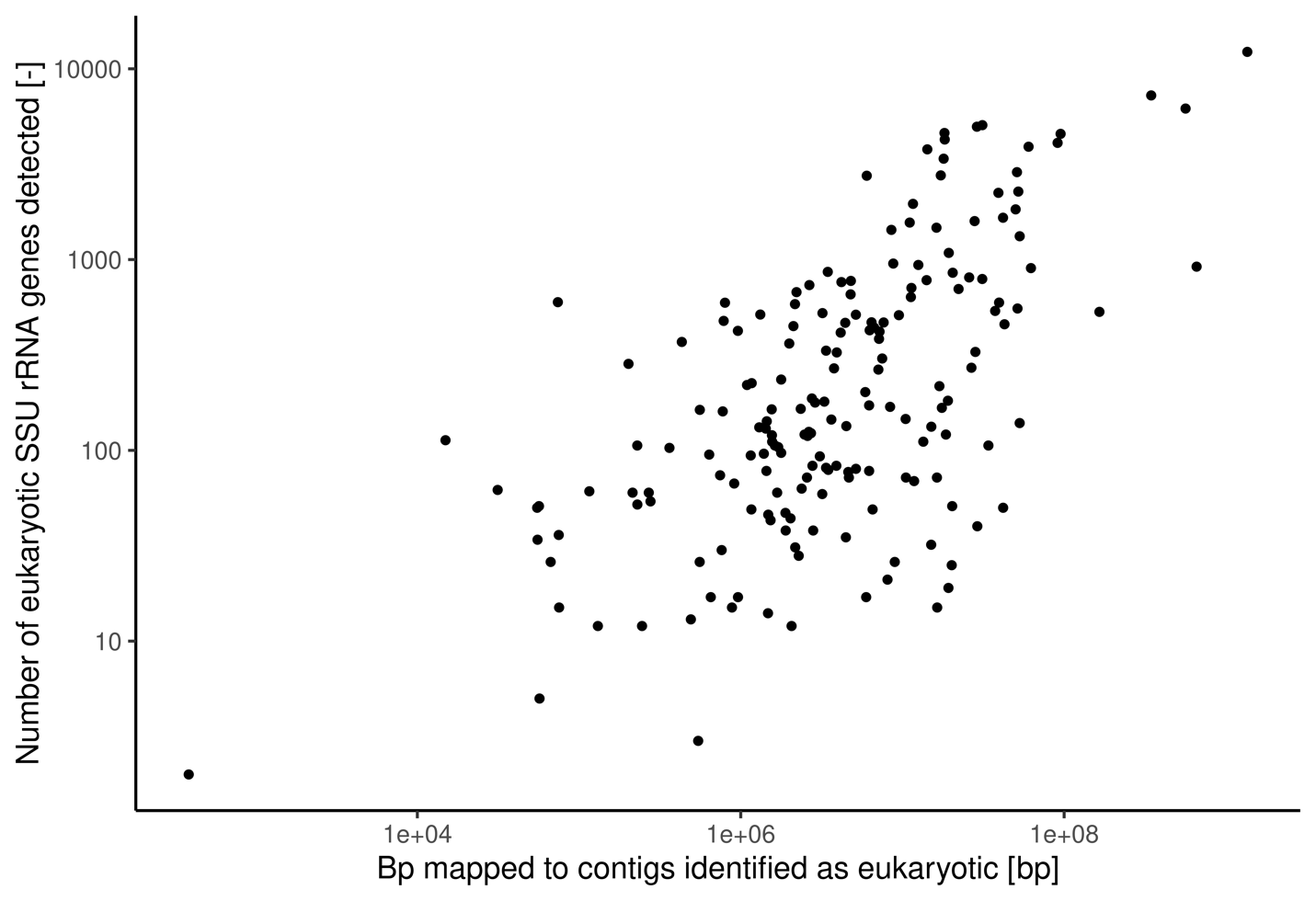

Figure S6. Number of eukaryotic SSU rRNA genes identified from the cleaned reads compared to the bp mapped to the contigs identified as eukaryotic. Spearman correlation coefficient = 0.58 (p-value < 0.001)

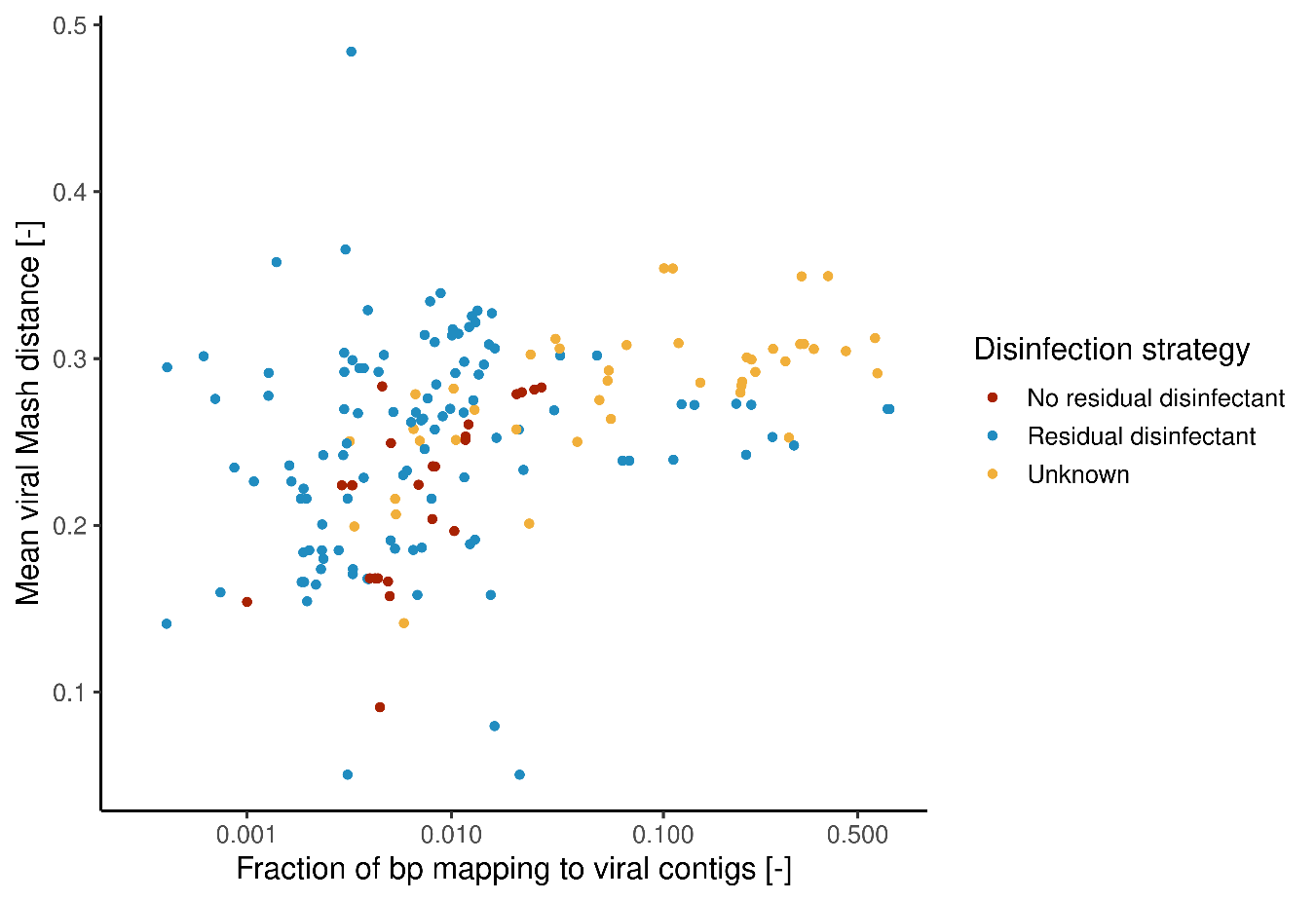

Figure S7. Viral Mash distance as a function of the prokaryotic fraction of the metagenomes investigated (EUKsemble results refined based on Kaiju’s taxonomic classification). Spearman correl. disinfected systems: 0.18 (p-value: 0.09); non-disinfected systems: 0.69 (p-value: < 0.001).

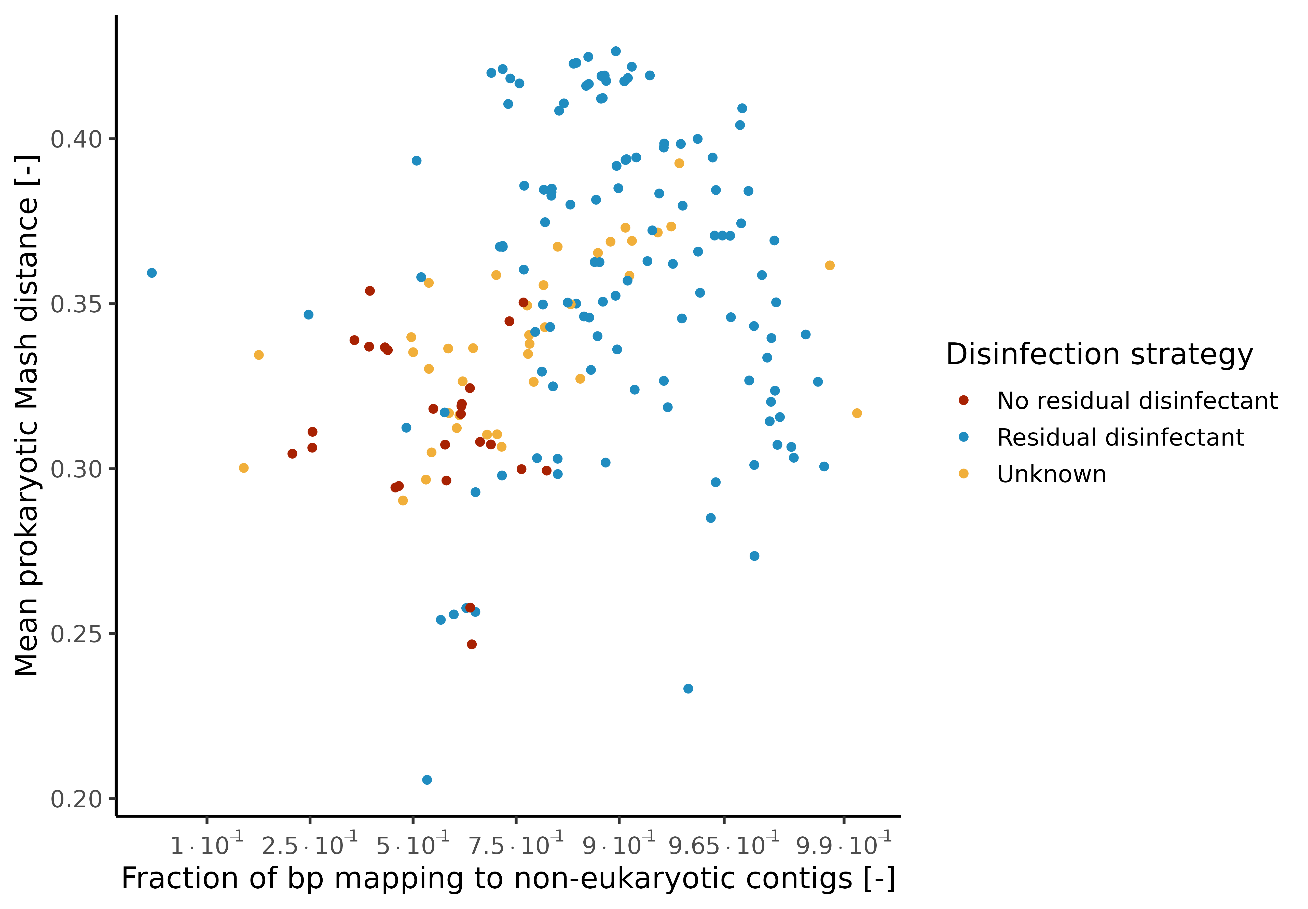

Figure S8. Prokaryotic Mash distance as a function of the prokaryotic fraction of the metagenomes investigated (EUKsemble results refined based on Kaiju’s taxonomic classification). Spearman correl. disinfected systems: 0.019 (p-value: 0.85); non-disinfected systems: -0.13 (p-value: 0.752).
